# Tempo and mode of genome evolution in a 50,000-generation experiment

**DOI:** 10.1101/036806

**Authors:** Olivier Tenaillon, Jeffrey E. Barrick, Noah Ribeck, Daniel E. Deatherage, Jeffrey L. Blanchard, Aurko Dasgupta, Gabriel C. Wu, Sébastien Wielgoss, Stéphane Cruveiller, Claudine Médigue, Dominique Schneider, Richard E. Lenski

## Abstract

Adaptation depends on the rates, effects, and interactions of many mutations. We analyzed 264 genomes from 12 *Escherichia coli* populations to characterize their dynamics over 50,0 generations. The trajectories for genome evolution in populations that retained the ancestral mutation rate fit a model where most fixed mutations are beneficial, the fraction of beneficial mutations declines as fitness rises, and neutral mutations accumulate at a constant rate. We also compared these populations to lines evolved under a mutation-accumulation regime that minimizes selection. Nonsynonymous mutations, intergenic mutations, insertions, and deletions are overrepresented in the long-term populations, supporting the inference that most fixed mutations are favored by selection. These results illuminate the shifting balance of forces that govern genome evolution in populations adapting to a new environment.

---

Adaptation to a new environment is a complex process that depends on the rates, effects, and interactions of mutations. Comparative genomics has revealed the molecular basis of some specific adaptations including such examples as lactase permanence in humans (*1*), domestication of plants (*2*) and animals (3), and pathogenicity in bacteria (4). Nevertheless, it has been difficult to determine more broadly what fraction of new mutations appearing in an evolving lineage are beneficial versus neutral or deleterious. Answering this question is important for modeling changes in sequences that are the basis of phylogenetic methods (5) and would inform debate about the relative importance of adaptive versus nonadaptive modes of genome evolution (*6, 7*).

The combination of experimental evolution and whole-genome sequencing provides a way forward owing to controlled conditions, replicate populations, and temporal data (8). In one study, the diversity of mutations involved in the early stage of adaptation to high-temperature stress was studied by sequencing >100 bacterial lineages after a 2000- generation experiment (9). In another study, sequencing a series of clones from one population over 40,000 generations showed the trajectory of genome evolution (10). However, these studies had some limitations: a short-term experiment reveals only the early steps of adaptation, and it is difficult to distinguish between adaptive “driver” and nonadaptive “passenger” mutations when only one population is examined.

To overcome these limitations, we analyzed the complete genomes of 264 *Escherichia coli* clones sampled from 12 populations across 50,000 generations of the longterm evolution experiment (LTEE) (*11, 12*). These populations have been propagated in a defined medium with scarce resources since 1988. The mean fitness of the bacteria, measured in competition with their ancestor, has increased by ∽70% over that time (12). The LTEE is a model system for studying fundamental evolutionary questions (*10-19*).

## Genome-wide mutations and hypermutability

We sequenced the genomes of 2 clones from each population after 500, 1000, 1500, 2000, 5000, 10,000, 15,000, 20,000, 30,000, 40,000 and 50,000 generations (Table S1) using the Illumina platform, and we predicted the mutations in each genome, including structural variation, using the *breseq* pipeline (20, 21). In total, we found 14,572 point mutations; 500 IS-element insertions; 726 deletions and 1132 insertions each ≤50 bp (small indels); and 267 deletions and 45 duplications each >50 bp (large indels). After 50,000 generations, the average genome length declined by 63 kbp (∽1.4%) relative to the ancestor (Fig. S1). Mutations were not distributed uniformly across the populations. Instead, six populations (Ara-1, Ara-2, Ara-3, Ara-4, Ara+3 and Ara+6) had 96.5% of the point mutations, having evolved hypermutable phenotypes caused by mutations that affect DNA repair or the removal of oxidized nucleotides (13-15). Fig. 1A shows the trajectories for the total number of mutations in all 12 populations; Fig. 1B is rescaled to provide better resolution for those that did not become point-mutation mutators.

**Fig 1.**
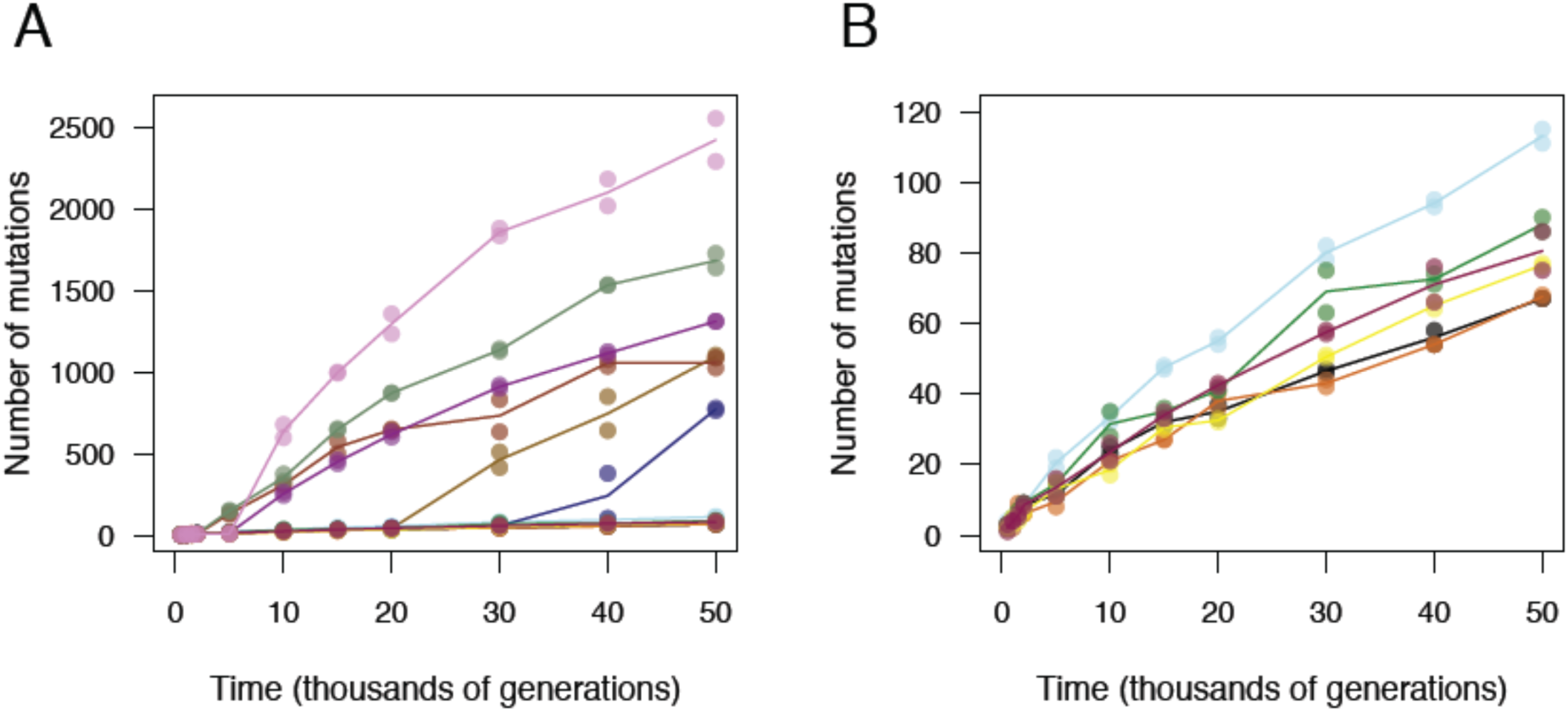
Total number of mutations over time in the 12 LTEE populations. **(A)** Total mutations in each population. **(B)** Total mutations rescaled to reveal the trajectories for the six populations that did not become hypermutable for point mutations. Each colored symbol shows a sequenced genome; each colored line passes through the average of the two genomes from the same population and generation.

Increased numbers of IS elements can also cause hypermutability (22), with higher rates not only of transpositions but also deletions and duplications through homologous recombination. In population Ara+1, 31.8% of all mutations observed through 50,000 generations were IS150 insertions, compared with 12.3% for the other five populations that also never evolved an elevated rate of point mutations. This mode of hypermutability evolved early in Ara+1; IS150 insertions are overrepresented in each Ara+1 clone from 5,000 generations onward when compared individually to all other non-point-mutator clones from the same generation (Fisher’s exact test with Bonferroni correction, *p* < 0.05). Two clones from other populations were also IS150 hypermutators by this test: 38.7% of the mutations in one 30,000-generation clone from Ara-5 and 31.7% of the mutations in one 40,000-generation clone from Ara-3 were IS150 insertions. The aberrant Ara-5 clone shares only one early mutation with other sequenced Ara-5 clones, indicating that it represents a deeply diverged lineage in that population; furthermore, it does not share point mutations with any other population, excluding cross-contamination. The emergence of these various mutator types shows how evolution can alter the production of genetic diversity (*15, 23*), which in turn will change the tempo and mode of genome evolution.

## Population phylogenies

Fig. 2A shows phylogenetic trees constructed using point mutations for each population; Fig. 2B shows the trees with branches rescaled after the point-mutation mutators evolved. Some populations—including Ara-2, which became hypermutable early, and Ara-6, which never did—harbor lineages that coexisted for tens of thousands of generations. Some other populations—such as Ara-4, which became hypermutable, and Ara+2, which did not—are more linear in structure, without deep branches among the sequenced clones. The deepest branches were likely supported by the diversity-promoting effects of negative frequency-dependent interactions, as demonstrated in the Ara-2 population (*18, 19*).

**Fig 2.**
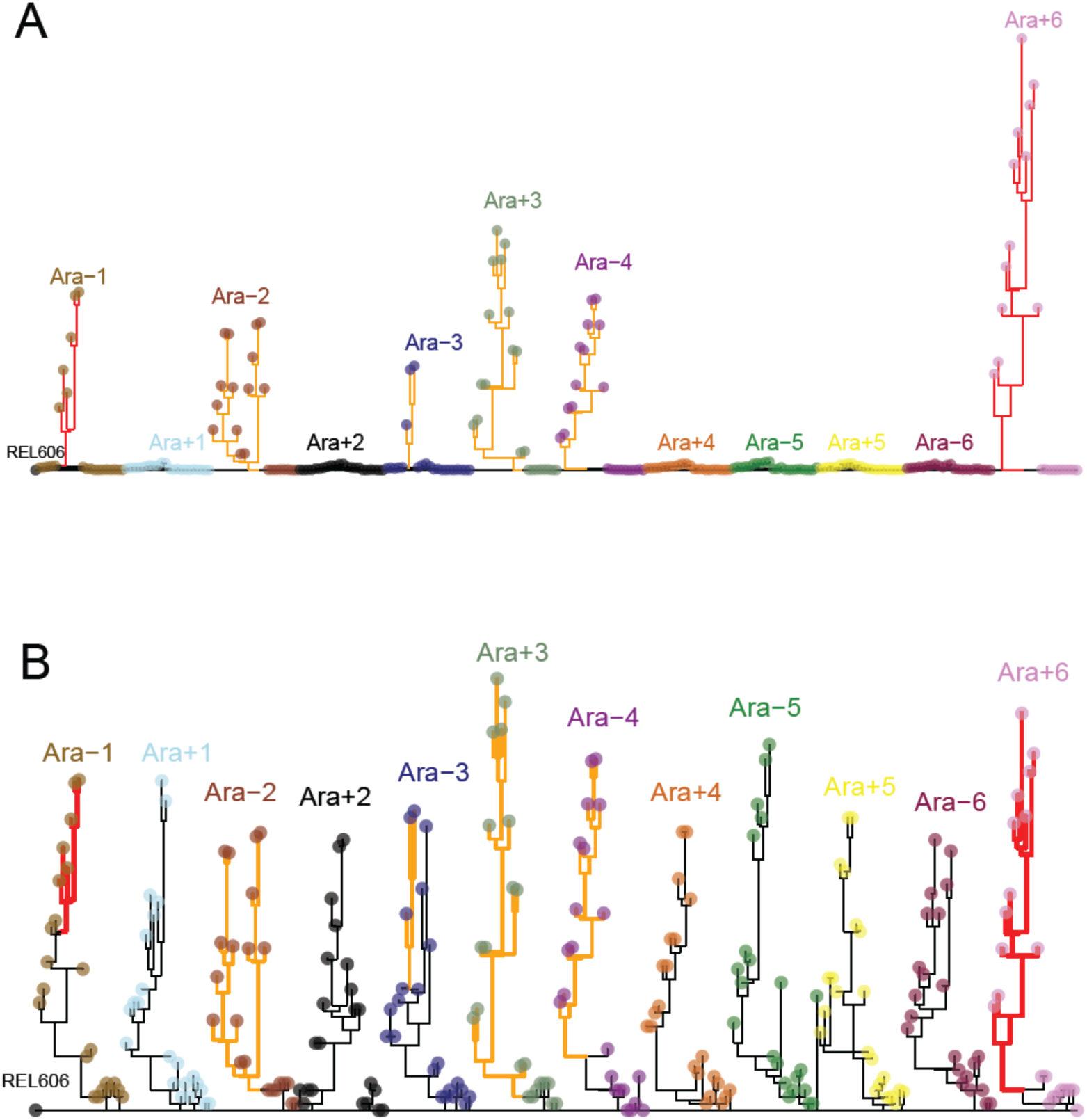
Phylogenetic trees for LTEE populations. **(A)** Phylogenies for 22 sequenced genomes sampled from each population, based on point mutations. The color scheme used here for the 12 populations is employed in other figures, when relevant. **(B)** The same trees, except branches after the evolution of mutators are rescaled to show their structure more clearly. Branches for lineages with mismatch-repair defects are orange and shortened by a factor of 25; branches for mutT mutators are red and shortened by a factor of 50. Strain REL606 (at left) is the ancestor of the LTEE. No early point mutations are shared between any populations, confirming independent evolution. Most populations have multiple basal lineages that reflect early diversification and extinction; some have deeply divergent lineages with sustained persistence, most notably Ara-2.

## Dynamics of genome evolution

The genome-wide substitution rate for point mutations increased by ∽100-fold in several populations after mutations arose that caused hypermutability (*13-15*), and these changes overwhelm the genomic signature of adaptation. Although the mutator lineages got a slight boost in their rate of fitness gain, the effect is small owing to clonal interference between beneficial mutations (*12, 24*). As a consequence, beneficial mutations are lost in a sea of unselected mutations in mutator populations. Therefore, to better understand the dynamic coupling between adaptation and genome evolution, we analyzed only the populations that retained the ancestral mutation rate (*25)* through 50,000 generations and the others before they became point-mutation or *IS150* mutators.

Wiser *et al*. (12) showed the LTEE’s trajectory for mean fitness is well described by a power-law relation, in which log fitness increases linearly with log time. Moreover, the power law accurately predicts fitness out to 50,000 generations using data from the first 5,000 generations only. Wiser *et al*. also showed that a population-dynamical model that incorporates two phenomena known to be important in the LTEE—clonal interference (*8, 26*) and diminishing-returns epistasis (*8, 17*)—generates the power-law relationship. This dynamical model further predicts that the number of beneficial mutations in a lineage increases with the square root of time (12). However, not all mutations that accumulate are beneficial; neutral and nearly neutral mutations can spread by recurring mutation, by drift, and by hitchhiking with beneficial mutations. Selective sweeps will purge some neutral mutations while causing others to increase; the expected overall effect is that the average number of neutral mutations in a lineage should increase linearly with time (*15, 27*).

To test these predictions, we fit three models to the trajectory for the number of mutations that accumulated over time in the nonmutator and premutator lineages:

*m* = *at*

*m* = *b* sqrt(t)

*m* = *at* + *b* sqrt(t)

where *m* is the number of mutations, *t* is time (generations), and *a* and *b* govern the genome-wide rates of accumulation of neutral and beneficial mutations, respectively (Fig. 3). (Fig. S2 shows the models fit to each population separately.) We compared the models using the Akaike information criterion (AIC), and the two-parameter model fits the data much better than those with only the linear (ΔAIC = −77.7) or square-root (AIC = −99.7) terms. Because the one-parameter models are nested within the two-parameter model, we can also assess the significance of adding the second parameter to the model; the resulting p-values for adding the parameter are 7.5 x 10'^-5^ and 5.2 x 10'^-7^ relative to the linear and square-root models, respectively. The trajectory for genome evolution thus shows strong signatures of both adaptive and nonadaptive changes.

**Fig 3.**
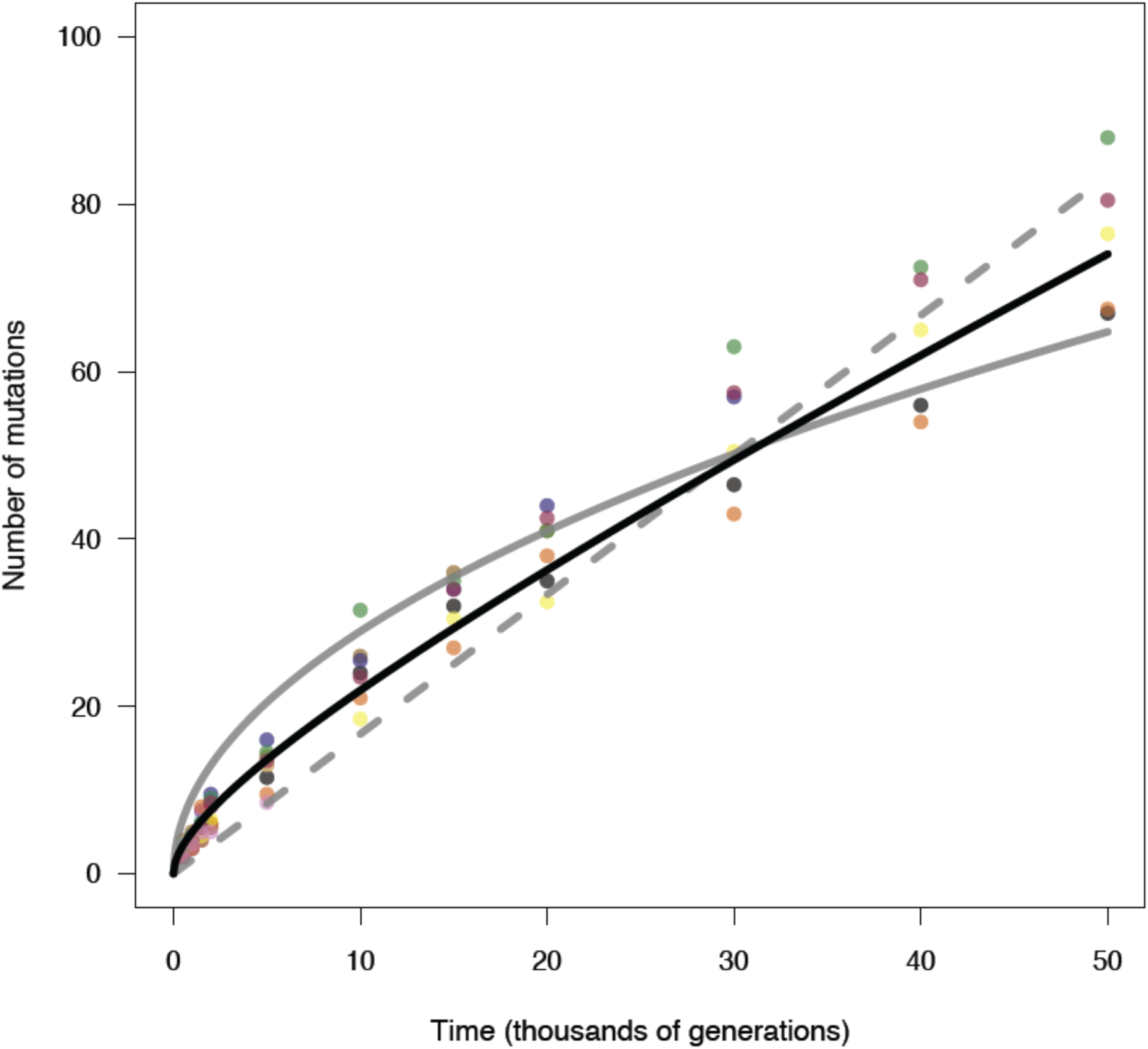
Alternative models fit to trajectory of genome evolution. Each symbol shows the total number of mutations for one LTEE population, based on the mean of its sequenced genomes at that time. Data are shown for populations that never became mutators and for other populations before hypermutability evolved. The dashed grey line shows the best fit to the linear model, *m* = *at*. The solid grey curve shows the best fit to the square-root model, *m* = *b* sqrt(t). The solid black curve shows the best fit to the composite model, *m* = *at* + *b* sqrt(t), where *a* = 0.000944 and *b* = 0.134856. See text for statistical analysis.

## Most observed mutations are beneficial

What proportion of the genomic changes in the nonmutator populations was adaptive, and how did it change over time? Fig. 4A shows the cumulative fraction of beneficial mutations estimated using the two-parameter model above, which declined from 86.3% at 500 generations to 38.7% at 50,000 generations. Fig. 4B shows the instantaneous proportion of adaptive changes, which dropped to 23.9% at 50,000 generations. Figs. 4C and 4D show the same estimates as the ratio of total mutations to the expectation under the null model where all mutations are neutral; a ratio of 2, for example, indicates 50% beneficial mutations, and 1 means all mutations are neutral. This latter representation is useful for showing two other approaches.

**Fig 4.**
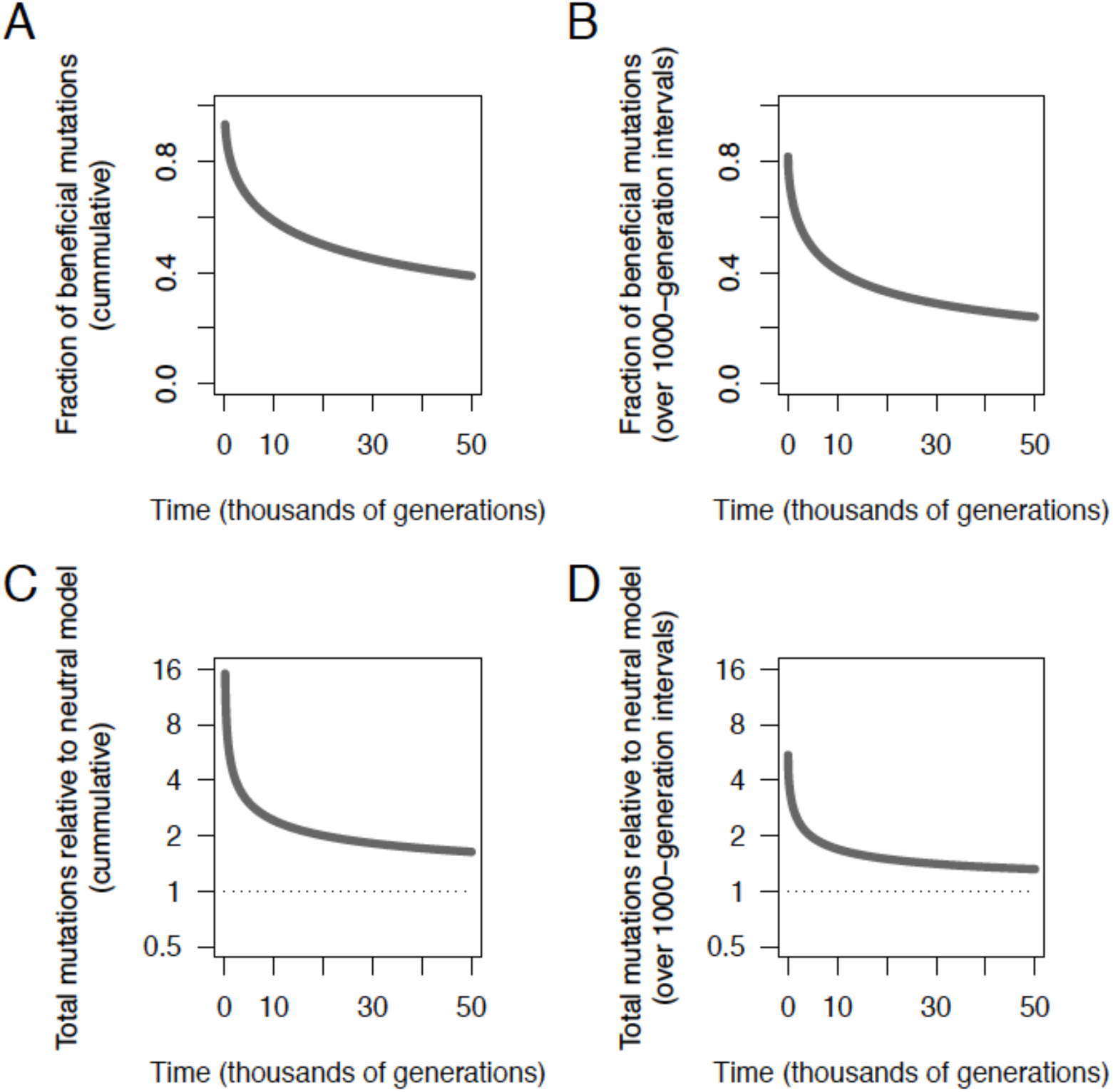
Contribution of beneficial mutations to genome evolution. **(A)** Cumulative fraction of genome changes resulting from beneficial mutations, based on best fit of the composite model in Fig. 3. (**B**) Fraction of changes from beneficial mutations over successive 1000-generation intervals. (**C**) Total mutations relative to the expectation under the null model where all mutations are neutral. (**D**) Total mutations relative to the neutral model over successive 1000-generation intervals.

A second approach uses the genomic data more directly, but only point mutations; it reflects the expectation that synonymous substitutions—point mutations in protein-coding genes that do not affect the amino-acid sequence—are neutral and should accumulate at a rate that equals their underlying mutation rate (*25, 27*). This expectation is not strictly true owing to selection on codon usage, RNA folding, and other effects, but it is generally thought that selection on synonymous changes is extremely weak, affects only a small fraction of sites at risk for synonymous mutations, or both (28). We calculate whether nonsynonymous and intergenic point mutations are found in excess relative to synonymous mutations, given the number of sites at risk for each class of mutation. Fig. 5A shows the number of synonymous mutations in nonmutator and premutator populations, scaled such that the mean at 50,000 generations is unity. As expected, synonymous mutations accumulated at an approximately constant rate (Fig. S3). Fig. 5B shows the number of nonsynonymous mutations relative to the neutral expectation based on synonymous mutations. Nonsynonymous mutations accumulated ∽17.1 times faster than synonymous ones over the first 500 generations and ∽3.4 times faster over the entire 50,000 generations. The cumulative 3.4:1 ratio implies that ∽70% (i.e., 2.4/3.4) of the nonsynonymous mutations observed in nonmutator lineages were beneficial—that is, they increased in frequency owing to positive selection. Nonsynonymous mutations still accumulated at over twice the rate of synonymous mutations in later generations (Fig. S4), implying that most nonsynonymous mutations were beneficial even after tens of thousands of generations in a constant environment. The same approach applied to intergenic point mutations (Fig. 5C) also reveals a large excess relative to synonymous mutations (6.5:1), implying that ∽85% of observed intergenic point mutations were adaptive.

**Fig 5.**
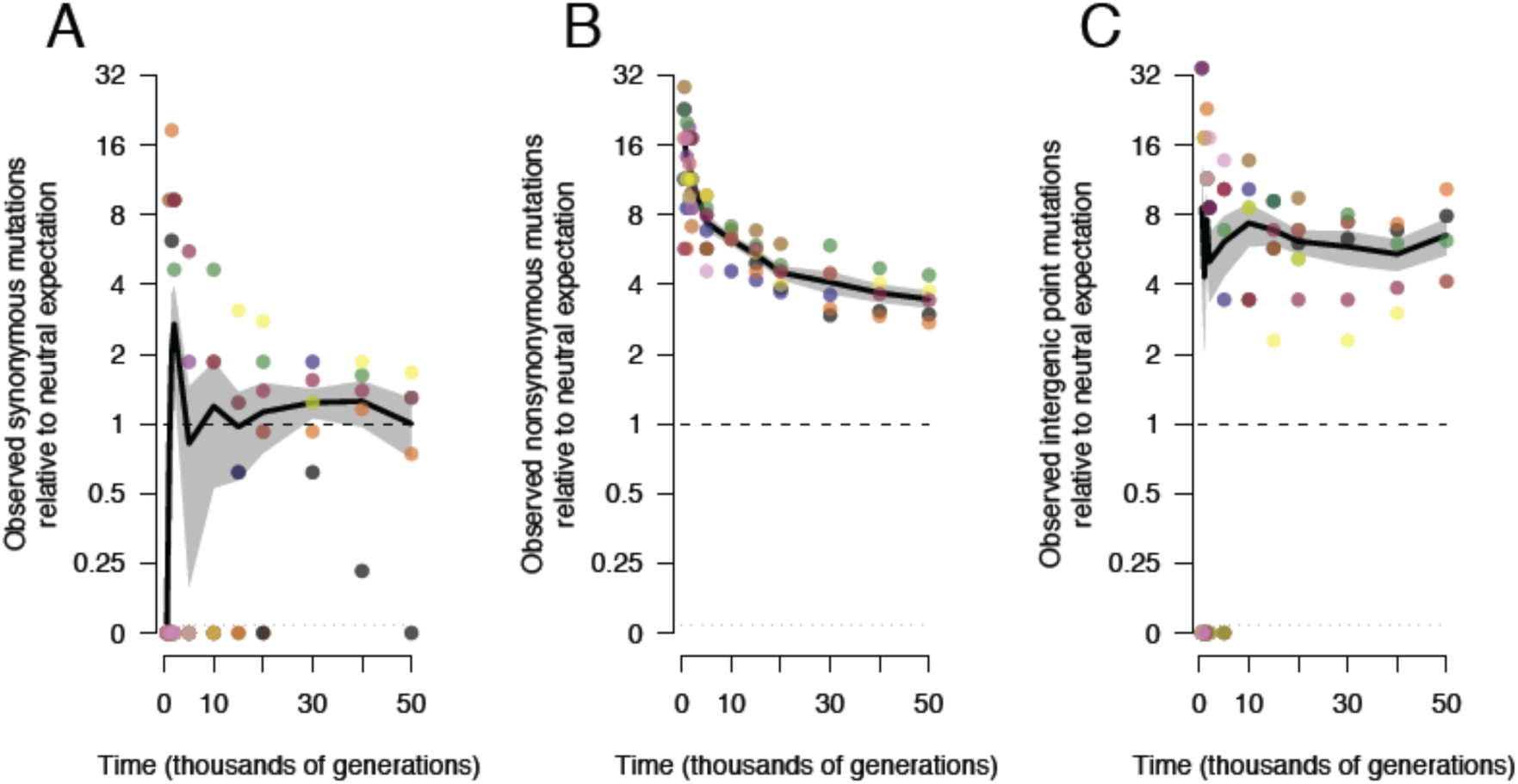
Trajectories for synonymous, nonsynonymous, and intergenic point mutations. **(A)** Accumulation of synonymous mutations, scaled such that the mean of the five nonmutator populations at 50,000 generations is unity. **(B)** Accumulation of nonsynonymous mutations, scaled using the same rate as synonymous mutations after adjusting for the number of genomic sites at risk for nonsynonymous and synonymous mutations. **(C)** Accumulation of intergenic point mutations, scaled using the same rate as synonymous mutations after adjusting for the sites at risk for intergenic and synonymous mutations. Each symbol shows the mean of the sequenced genomes from the nonmutator or premutator lineages in one population. Note the discontinuous scale, in which populations with no mutations of the indicated type are plotted below. Black lines connect grand means; the grey shading shows standard errors calculated from the replicate populations.

Synonymous mutations provide an internal benchmark that reveals the role of positive selection in the accumulation of nonsynonymous and intergenic point mutations. However, synonymous mutations are not directly informative for understanding how selection affects the accumulation of insertions and deletions that collectively comprise 45.6% of all mutations in the nonmutator and premutator clones sequenced through 50,000 generations. This set of mutations includes 54.4% base substitutions, 23.2% IS insertions, 8.7% small indels (≤50 bp), 10.6% large deletions (>50 bp), and 3.0% large duplications (>50 bp). Despite changes in the rate at which mutations accumulated (Fig. 3) and the fraction of beneficial mutations (Fig. 4), the composition of observed mutations with respect to these categories was relatively stable over time (Fig. S5). Again considering only nonmutator clones, there was no significant difference in the types of mutations that arose before 10,000 generations and those present in the 50,000-generation clones that were not in the 40,000-generation clones (Fisher’s exact test, *p* = 0.53). This constancy in the spectrum of changes over time implies that no one type of mutation was much more likely to generate a benefit than other types, and it suggests that the fraction of beneficial mutations for other mutation types is roughly similar to that for base substitutions.

We developed a third approach to estimate the proportion of beneficial changes that can be applied to all types of mutation. We compare the mutational spectrum between two experiments, the LTEE and a Mutation Accumulation Experiment (MAE) in which 15 lines were propagated via single-cell bottlenecks (*29*). Such bottlenecks eliminate the genetic variation needed for natural selection, so that all types of mutations accumulate at the rates at which they happen spontaneously, regardless of their fitness effects, except for lethal or severely deleterious mutations that preclude cells from making colonies used to propagate the lines (8). Such unselected lines thus provide a baseline for distinguishing beneficial and nonbeneficial mutations. In fact, because more unselected mutations are deleterious than beneficial, MAE lines are expected to lose fitness over time, and they did (Fig. S6).

To quantify the relative rates for all types of mutations in the absence of selection, we sequenced clones from the MAE lines after 550 days of serial bottlenecks (Table S1). Consistent with the random accumulation of mutations, the number of nonsynonymous mutations (including nonsense) was similar to the expectation based on synonymous mutations (117 observed, 105.02 expected); the resulting ratio of 1.11 is well within the 95% confidence interval (0.70-1.50) obtained by a randomization test. Also, there was no among-line variation in the total number of mutations (*χ*^2^ = 5.46, df = 14, *p* = 0.978). We can therefore reasonably use the MAE lines to estimate the relative rates of different types of mutations, with synonymous mutations providing a benchmark that should be largely free of selection in both experiments. For example, population Ara-1 had 21 nonsynonymous mutations at 20,000 generations and the expected number of synonymous mutations based on the average nonmutator LTEE population was 1.08 (Fig. S3); the 15 MAE lines in total had 117 nonsynonymous and 39 synonymous mutations; thus, the ratio of observed mutations to the neutral expectation in this case is (21/1.08)/(117/39) = 6.5. These ratios show that all major classes of mutations—including various insertions and deletions—are substantially overrepresented in the LTEE relative to the MAE (Fig. S7), implying that many mutations in each class were adaptive during the LTEE.

All three approaches, based on different assumptions and comparisons, indicate that most mutations observed in the nonmutator LTEE populations were beneficial drivers rather than mere passengers. Also, the proportion of observed mutations that were beneficial declined over time but remained high throughout the LTEE’s 50,000 generations.

## Parallel evolution at many gene loci

Parallel evolution occurs when similar changes arise independently in multiple lineages at a frequency above a suitable null model, and it is often used to discover putative targets of selection (*4, 9, 16, 30*). Genetic parallelism can be studied at the level of DNA sequence, affected genes, or integrated functions. Parallelism at the nucleotide level tends to be rare because different mutations in a gene often produce similar benefits (*4, 9, 16*), although there are exceptions (*30*). Parallelism at a functional level requires detailed understanding that is often unavailable, and it is difficult to interpret when there are many mutations. We therefore examined parallelism at the gene level.

We focused on lineages that retained the ancestral point-mutation rate (including clones from populations that later became hypermutable) because, as shown above, most mutations are drivers in those cases; hypermutability makes such analyses less informative because many more mutations are passengers. We first calculated the expected number of nonsynonymous mutations for each single-copy protein-coding gene based on its length as a fraction of all such genes and the total number of nonsynonymous mutations in the relevant lineages (Table S2). We computed *G* scores for goodness-of-fit between observed and expected values; the total score was 2592.9. We compared that total with simulated datasets where positions of mutations in the coding genome were randomized, and the observed total significantly exceeded the simulations (mean simulated *G* = 1933.7, *Z* = 25.5, *p* < 10^-144^). Fifty-seven genes had two or more mutations; these genes had 50.1% of the nonsynonymous mutations but constituted only 2.1% of the coding genome. (In contrast, only one gene had multiple synonymous changes.) Table 1 shows the 15 genes that contribute the most to the total *G* score for nonsynonymous mutations. Several encode proteins with core metabolic or regulatory functions, including three involved with peptidoglycan synthesis.

**Table 1.**
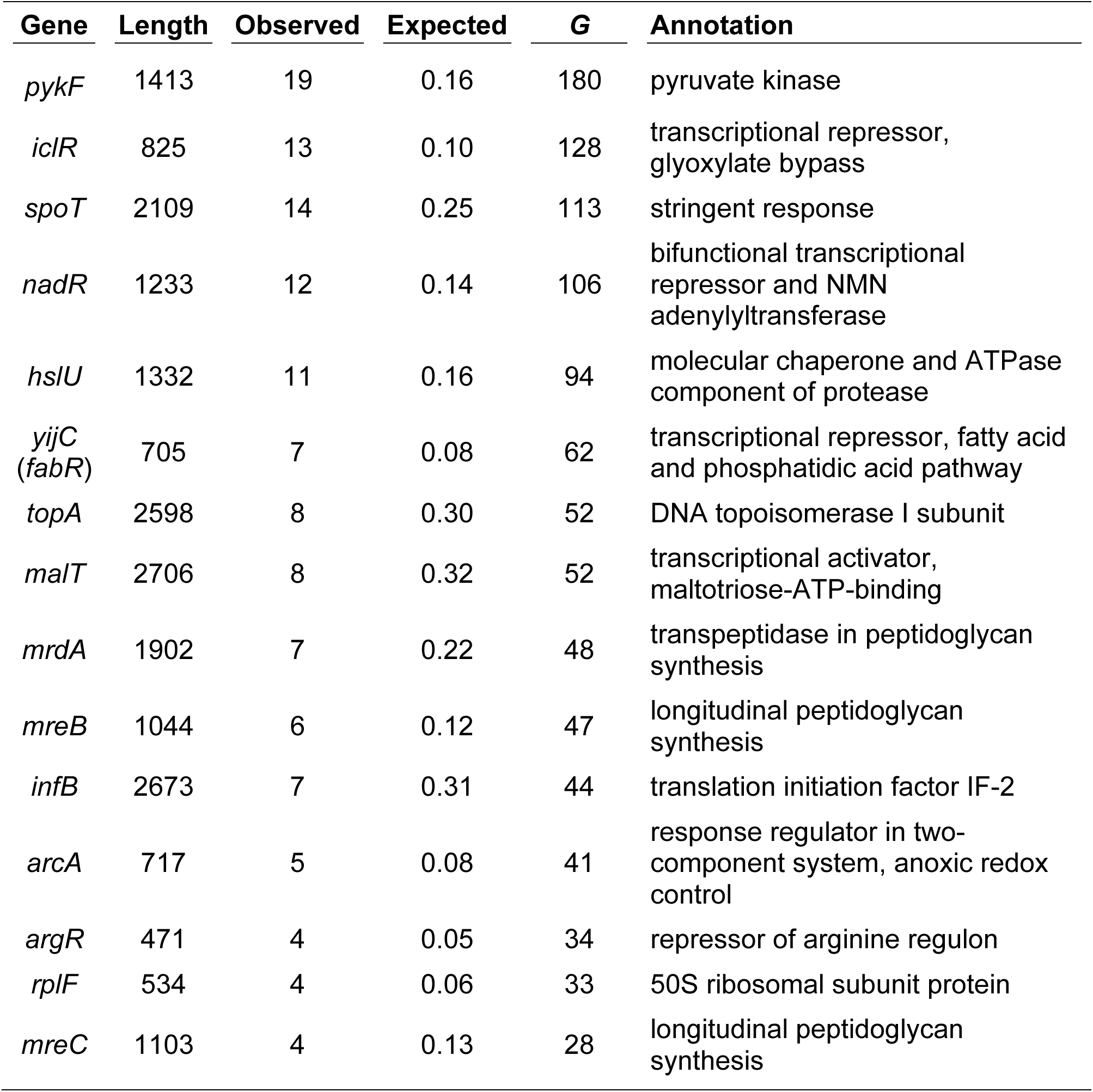
Protein-coding genes with highest *G* scores. Genes are ranked by *G* scores computed using the observed number of independent nonsynonymous mutations relative to the number expected given the gene length (bp). The parenthetical gene name is a synonym. The data are from populations that retained the ancestral point-mutation rate throughout the 50,000 generations and from other populations before they evolved hypermutability.

Table 2 lists the 16 genes with the most deletions, duplications, insertions, and intergenic point mutations. In 12 cases, the majority of the mutations were mediated by IS elements; these include new insertions as well as deletions and duplications involving preexisting IS elements that can facilitate such mutations via homologous recombination. In six cases (including five involving IS-element insertions), the same or nearly identical mutations occurred in one or more of the MAE lines, where the opportunity for selection was minimal, suggesting that these cases might involve mutational hotspots. While parallel changes involving IS elements may indicate high-frequency mutational events, recall that IS insertions and large indels are substantially enriched in the LTEE populations relative to the MAE lines (Fig. S7), implying that many of those mutations are also beneficial. Indeed, the IS-mediated *rbsD* deletions have been shown both to occur at a high rate and to confer an advantage in the LTEE environment (*31*), and some IS-mediated mutations have been shown to be beneficial in other studies as well (*32, 33*).

**Table 2.** Genes with the most deletions, duplications, insertions, and intergenic point mutations. Genes are ranked based on total number of mutations excluding nonsynonymous and synonymous point mutations. When two genes are separated by a slash, the affected sequence includes the intergenic region between them. Parenthetical gene names are synonyms. The column labeled IS indicates whether the majority of mutations involve IS elements. The MAE column indicates whether the same or nearly identical mutations occurred in one or more of the MAE lines. The data are from populations that kept the ancestral point-mutation rate throughout the 50,000 generations and other populations before they evolved hypermutability.

| Genes | Mutations | Number | IS | MAE | Annotation |
| --- | --- | --- | --- | --- | --- |
| *rbsD* | mostly large deletions | 41 | yes | no | D-ribose utilization; most deletions affect entire *rbs* operon |
| *nupC* | various intergenic | 19 | yes | yes | nucleoside transporter |
| *iap* | mostly large indels | 19 | yes | no | alkaline-phosphatase isozyme conversion; most indels affect tens of adjacent genes including *rpoS*, which encodes stationary-phase $\sigma$ factor |
| *mokB* | various indels | 17 | yes | yes | enables *hokB* toxin expression |
| *yhgl/gntT* | intergenic point mutations | 16 | no | no | gluconate transport |
| *mokC* | various indels | 15 | yes | yes | enables *hokC* toxin expression |
| *ybcU (borD)* | large indels | 14 | yes | no | indels affect this and adjacent remnants of DLP12 prophage |
| ECB_02013 | various indels | 14 | no | yes | indels affect this and adjacent remnants of P2-like prophage |
| ECB_02816 (*kpsD*) | various indels | 14 | yes | no | polysialic-acid transport protein precursor |
| *acs/nrfA* | various intergenic | 14 | no | no | acetyl-CoA synthase; nitrite reductase |
| *hokE* | large indels | 12 | yes | no | toxin in plasmid-derived toxin-antitoxin system; most indels affect several adjacent genes involved in iron acquisition |
| *ybeB/phpB* | various intergenic | 11 | yes | no | unknown functions, but adjacent to genes involved in cell-wall synthesis |
| *ydiJ/ydiK* | various intergenic | 11 | no | no | predicted FAD-linked oxidoreductase; putative inner membrane protein |
| *ldrC* | various indels | 10 | yes | yes | small toxic polypeptide |
| *menC* | IS insertions | 10 | yes | yes | menaquinone biosynthesis |
| *fimA* | mostly IS insertions | 10 | yes | no | component of fimbrial complex |

The parallelisms involving nonsynonymous substitutions and other mutation types in the LTEE populations, coupled with their high rates of accumulation relative to the MAE lines, provide evidence that many mutations in the sequenced genomes were drivers of adaptation. For many insertions and deletions, however, the specific genes that are the foci of adaptive evolution are difficult to identify owing to the multiplicity of genes affected and the potentially confounding effect of mutational hotspots.

## Discussion

Adaptation by natural selection sits at the heart of phenotypic evolution. However, the random processes of spontaneous mutation and genetic drift often overwhelm, or at least obscure, the genomic signatures of adaptation. We overcame this difficulty by combining a well-controlled experimental system with whole-genome sequencing. In particular, we analyzed 264 genomes from 12 populations of *E. coli* that evolved for 50,000 generations under identical culture conditions. Even so, six of these populations evolved hypermutable phenotypes that caused their point-mutation rates to increase by ∽100-fold, and another evolved hypermutability mediated by a mobile genetic element. By focusing our analyses on populations that retained the ancestral mutation rate, we identified several key features of the tempo and mode of their genome evolution. First, a population-genetic model with two terms—one for beneficial drivers, the other for hitchhikers—describes the dynamics of genome change much more accurately than models without both terms. Second, the great majority of mutations in the genomes sampled during the early generations were beneficial drivers. Third, the proportion of observed mutations that were beneficial declined over time but remained substantial even after 50,000 generations. The second and third findings follow from the population-genetic model. Both are also strongly supported by the excess of nonsynonymous and intergenic point mutations relative to synonymous changes in the LTEE and by the excess of all major classes of mutations, including insertions and deletions, in comparison to the mutation-accumulation lines. Fourth, there was extensive gene-level parallel evolution involving all types of mutations across the replicate LTEE populations.

Our analyses also show a striking contrast between the contributions of beneficial mutations to molecular evolution and to the marginal fitness improvements observed after 50,000 generations in a stable environment. In particular, our data indicate that beneficial mutations continue to constitute a large fraction of genetic changes, whereas the fitness gains amount to only a few percent over the last 10,000 generations. Beneficial mutations with very small selection coefficients—including those well below our ability to measure them directly—are nonetheless visible to natural selection (12). Hence, adaptation remains a major driver of molecular evolution even many tens of thousands of generations after an environmental shift. Our experimental results thus support a selectionist view of molecular evolution, complementing indirect evidence based on comparative genomics in bacteria, *Drosophila,* and humans (*34-36*). Also, if the process of adaptation is still so pervasive such a long time after populations encounter new conditions, then fixations of neutral mutations will occur primarily by hitchhiking, or “genetic draft” (37-39), rather than by genetic drift as traditionally envisioned. Of course, the LTEE has some features that differ from many or most natural populations including a low mutation rate, the absence of sex or horizontal gene transfer, and a stable environment. As seen in the lineages that became hypermutable, a high mutation rate reduces the relative contribution of adaptation to molecular evolution. The effects of horizontal gene transfer and variable environments on the dynamic coupling between genomic change and adaptive evolution should also be examined further. Longterm experiments with microorganisms provide opportunities for rigorous analyses of these issues as well.

## ACKNOWLEDGMENTS

O.T., J.E.B., D.S., and R.E.L. conceived the project; R.E.L. and J.L.B. provided strains; O.T., J.E.B., D.E.D., A.D., G.C.W., S.W., S.C., and C.M. analyzed genomes and generated other data; N.R. developed theory; R.E.L., O.T., and J.E.B. wrote the paper. All authors contributed to the intellectual development of the paper and approved the submitted version. We thank N. Hajela for technical assistance, R. Maddamsetti and Z. D. Blount for discussions, and M. Lynch for starting the MAE lines. This research was supported by the US National Science Foundation (DEB-1451740 to R.E.L.), the BEACON Center for the Study of Evolution in Action (DBI-0939454 to R.E.L.), the European Research Council (FP7 grant 310944 to O.T.), the European Union (FP7 grant 610427 to D.S.), the French National Funding Agency (ANR-08-GENM-023-001 to D.S., O.T., and C.M), and the US National Institutes of Health (R00- GM087550 to J.E.B.). D.E.D. was supported by a traineeship from the Cancer Prevention and Research Institute of Texas. The authors acknowledge the use of high-performance computing resources at the Texas Advanced Computing Center. The authors declare that they have no competing interests. Genome data have been deposited in the NCBI Sequence Read Archive (http://www.ncbi.nlm.nih.gov/sra). The *breseq* analysis pipeline is available at GitHub (http://github.com/barricklab/breseq). Other analysis scripts will be made available on publication at the Dryad Digital Repository.

## Supplementary Materials

Materials and Methods

Figs. S1 to S7

References (*40-48)*

Tables S1 to S3

